# Characterization and optimization of variability in a human colonic epithelium culture model

**DOI:** 10.1101/2023.09.22.559007

**Authors:** Colleen M. Pike, Bailey Zwarycz, Bryan E. McQueen, Mariana Castillo, Catherine Barron, Jeremy M. Morowitz, James A. Levi, Dhiral Phadke, Michele Balik-Meisner, Deepak Mav, Ruchir Shah, Danielle L. Cunningham Glasspoole, Ron Laetham, William Thelin, Maureen K. Bunger, Elizabeth M. Boazak

**Author notes:** Authors contributed equally (Colleen M. Pike, Bailey Zwarycz). **Correspondence address:** Elizabeth M Boazak, PhD Associate Director, R&D Altis Biosystems, Inc 6 Davis Drive, Durham, NC 27709. USA.

## Abstract

Animal models have historically been poor preclinical predictors of gastrointestinal (GI) directed therapeutic efficacy and drug-induced GI toxicity. Human stem and primary cell-derived culture systems are a major focus of efforts to create biologically relevant models that enhance preclinical predictive value of intestinal efficacy and toxicity. The inherent variability in stem-cell-based complex cultures makes development of useful models a challenge; the stochastic nature of stem-cell differentiation interferes with the ability to build and validate robust, reproducible assays that query drug responses and pharmacokinetics. In this study, we aimed to characterize and reduce potential sources of variability in a complex stem cell-derived intestinal epithelium model, termed RepliGut^®^ Planar, across cells from multiple human donors, cell lots, and passage numbers. Assessment criteria included barrier formation and integrity, gene expression, and cytokine responses. Gene expression and culture metric analyses revealed that controlling for stem/progenitor-cell passage number reduces variability and maximizes physiological relevance of the model. After optimizing passage number, donor-specific differences in cytokine responses were observed in a case study, suggesting biologic variability is observable in cell cultures derived from multiple human sources. Our findings highlight key considerations for designing assays that can be applied to additional primary-cell derived systems, as well as establish utility of the RepliGut^®^ Planar platform for robust development of human-predictive drug-response assays.

## Introduction

There is an urgent need for practical models capable of predicting human clinical outcomes with respect to drug pharmacology, toxicology, and disposition. While animals remain the most common model system to test efficacy and de-risk liabilities prior to initiating clinical trials, species differences limit their predictive capacity (Olson *et al*., 2000; Monticello *et al*., 2017). These species differences are particularly relevant in the gastrointestinal system where rodent models have very poor predictive power, most likely owing to innate differences in diet and microbiome (Valatas, Vakas and Kolios, 2013; DeVoss and Diehl, 2014; Ananthakrishnan, Kaplan and Ng, 2020). Immortal cell lines derived from human colonic carcinomas, such as the Caco-2 cell line, have been a powerful *in vitro* tool for studying underlying biology and predicting drug disposition in the intestine for decades. However, these models are not sufficient to recapitulate native colonic physiology due to genome instability, lack of cellular diversity, abnormal drug transport kinetics, and poor representation of intestinal disease mechanisms such as inflammation in cultured cells; therefore, these cell lines have poor clinical predictive power (Sambuy *et al*., 2005; Lennernäs, 2007; Press and Grandi, 2008; Sun *et al*., 2008; Larregieu and Benet, 2013; VanDussen *et al*., 2015). As a result, more physiologically relevant *in vitro* cell-based models of the human intestine have been a significant area of development in recent years (Sato *et al*., 2009; Breslin and O’Driscoll, 2013; Ahmad *et al*., 2014; Wang *et al*., 2014; Almeqdadi *et al*., 2019; Dutton *et al*., 2019; Yoo and Donowitz, 2019; Franco, Da Silva and Cristofoletti, 2021; Markus *et al*., 2021).

Although the ability to isolate, culture, and store primary cells directly from human tissue has changed the landscape of useful *in vitro* cell models, primary cell isolation from adult human intestine has proven to be particularly challenging (Grossmann *et al*., 2003). Unlike many other adult tissues, the gastrointestinal epithelium exhibits rapid turnover and relies on resident crypt stem cells to replenish the differentiated epithelium, which consists of at least five distinct cell lineages (Barker, van de Wetering and Clevers, 2008; Gracz and Magness, 2014; Rees *et al*., 2020). Post-differentiation, mature GI epithelial cells have only a 5-7 day lifespan before undergoing programmed cell death and detaching from the epithelium, making isolation and culture of fully differentiated cells difficult (Gibson *et al*., 1989). This challenge can be overcome by stimulating cultured crypt-resident stem cells with environmental cues such as growth factors, to either maintain a proliferative phenotype or induce terminal differentiation to produce a mature epithelium (Grossmann *et al*., 2003). When seeded onto an extracellular matrix, intestinal stem cells have been shown to differentiate and self-organize into spherical organoids with an enclosed center cavity that mimics the environment of the intestinal lumen (Sato *et al*., 2009; Ahmad *et al*., 2014).

Intestinal crypt-derived organoids are a popular tissue culture model because they offer several advantages over current animal and cell culture-based platforms. Organoids are derived from primary gut epithelial stem cells, reflect the cellular diversity of the native intestinal segments, and can be cultured from human patient biopsies (Sato *et al*., 2009; Gracz and Magness, 2014; Dutton *et al*., 2019; Yoo and Donowitz, 2019). Despite these advantages, conventional gut organoid technologies are limited by the inability to easily access the apical (or luminal) aspect of the cells—the cell surface where nutrients, microbiota, and drugs naturally interact with the gut epithelium. Organoids also exhibit heterogeneous morphologies and biological functions that are difficult to synchronize between organoid structures. Organoids also grow in 3-dimensional (3D) space embedded in Matrigel^®^, rendering downstream readouts such as high-content imaging difficult to achieve for labs without access to confocal microscopy (Almeqdadi *et al*., 2019). Moreover, these aspects of organoids make it challenging to model and measure integrity of the epithelial barrier, which underlies many GI adverse events and diseases such as inflammatory bowel disease IBD (Hill *et al*., 2017).

To overcome the limitations of 3D organoid cultures, the RepliGut^®^ Planar model was developed to replicate primary human colonic epithelium in a monolayer format. RepliGut^®^ Planar consists of cultured gut epithelial stem cells developed into a differentiated epithelium on a 2-dimensional (2D) scaffold in conventional Transwell^®^ inserts (Wang *et al*., 2017). The model uses culture conditions that enable intestinal stem cells to be isolated and expanded from human tissue and plated onto a proprietary membrane-supported biomimetic hydrogel to model the polarized colonic epithelial monolayer. RepliGut^®^ Planar is a dynamic model sequentially comprised of both undifferentiated stem cells and differentiated absorptive and secretory cell lineages of the colon. The 2D planar open-faced geometry of this platform allows access to the apical surface of the polarized epithelium for study of compound interactions, recapitulating *in vivo* interactions of oral medications with epithelial cells in the lumen of the colon. Exposure of the basal compartment in membrane-supported inserts allows modeling of interactions between intestinal cells and treatments such as inflammatory cytokines and systemic therapeutics in development.

While technology now allows for physiologically relevant platforms to be developed, practical implementation and use of these platforms has proved challenging (Criss *et al*., 2021), highlighting the need to understand key drivers of variability and how to control for them. The inherent variability in complex primary cell-derived models limits the ability to build reproducible assays to query effects of therapeutics. This variability also makes it difficult to assess the value of assays that utilize these informationally rich model systems. The focus of this study was to investigate factors most likely to contribute to variability in a primary stem cell-derived intestinal culture system, such as multiple human donors, cell manufacturing lots, and model lifecycle characteristics. Assessment metrics included culture dynamics, barrier formation and maintenance, and gene expression analysis across multiple human donors, cell lots and cell passage numbers. We identified cell passage number as a significant source of variability affecting gene expression profiles. Donor-to-donor variability was minimal when evaluating culture dynamics. However, assessment of four human donors revealed donor-dependent differences in responses to proinflammatory cytokines, tumor necrosis factor alpha (TNFα) and interferon gamma (IFNγ), with respect to transepithelial electrical resistance (TEER) kinetics, IC50s, and magnitude of lactate dehydrogenase (LDH) and chemokine release. Our findings highlight that when sources of variability are well controlled, donor-specific differences in response to inflammatory stimuli persist, which is reflective of inter-individual biologic variability. Clinically validated TNFα- and IFNγ antagonists, adalimumab and tofacitinib, mitigated cytokine-induced barrier disruption and cytotoxicity, demonstrating the utility of the RepliGut^®^ Planar as a tool to explore drug pharmacology associated with TNFα-and IFNγ pathways in inflammatory diseases. Together, these results identify key considerations for designing robust assays that can be applied to additional primary-cell derived systems, as well as establish utility of the RepliGut^®^ Planar platform for robust development of human-predictive drug-response assays.

## 2 Materials and Methods

### Cell Culture

Human intestinal tissue was obtained post-mortem via an established Organ Procurement Organization following consent of family under strict ethical guidelines established by the Organ Procurement Transplantation Network (OPTN; https://optn.transplant.hrsa.gov/). All donors tested negative for HIV I/II, Hepatitis B (HBcAB, HBsAG), and Hepatitis C (HCV) and were free of known intestinal diseases. Intestinal crypts were isolated from human transverse colon, expanded under sub-confluent conditions, and cryopreserved as described previously with minor modifications (Grossmann *et al*., 1998; Wang *et al*., 2017). Cell lots were defined as a single vialing and freeze-down process of cells pooled following expansion. For each experiment, vials were rapidly thawed in a 37°C water bath directly and seeded on hydrogel coated 12- or 96-well Transwell^®^ plates (Corning 3460 or 3392) at a density of 5-8 x 10^4^ cells/cm^2^ in RepliGut^®^ Growth Medium (RGM, Altis Biosystems, Durham, NC). Once cells reached confluence, media was changed to RepliGut^®^ Maturation Medium (RMM, Altis Biosystems, Durham, NC) to promote cellular differentiation and polarization. Media volumes were 1000 µl/2000 µl apical/basal for 12-well plates and 100 µl/200 µl apical/basal for 96-well plates. Media was changed every 48 hours except for the switch to RMM which was based on confluence without regard to timing of the previous media change. Viability was determined using a trypan blue exclusion assay on a Countess™ II automated cell counter (Invitrogen, AMQAX1000). Cytokine and cytokine inhibitor treatments were performed by adding treatments to both the apical and basal transwell compartments for 48 hours on day 2 in RMM. TNFα (R&D Systems, 210-TA) or IFNγ (PeproTech, 300-02), was added with or without simultaneous treatment with adalimumab (Selleck Chemicals A2010) or tofacitinib (Selleck Chemicals S2789), respectively.

### Transepithelial Electrical Resistance (TEER)

Barrier integrity of cell monolayers was assessed via TEER using an Epithelial Volt/Ohm Meter (World Precision Instruments, EVOM2 or EVOM3) and STX100C96 electrode for 96-well cultures or STX2 electrode for 12-well cultures. TEER was measured daily during experiments. Percent change in TEER of cytokine-treated samples (TNFα or IFNγ) from vehicle was calculated using the following equation: - ((Average TEER_Vehicle_ –TEER_Sample_)/Average TEER_Vehicle_)*100. IC_50_s were calculated using three-parameter nonlinear regression in Prism software (GraphPad Software, La Jolla, CA). Curve fitting was performed for all experimental runs, with an R-squared threshold of 0.6 applied as the minimum acceptable value and bounds on the 95% confidence interval of the computed IC_50_ within the range of doses tested. Runs that failed to achieve these quality metrics were deemed unsuccessful, primarily due to an inadequate dose range for a reliable curve fit and were consequently excluded from the reported TEER-based dose response metrics.

### Cell Fixation & Staining

For 5-Ethynyl-2′-deoxyuridine (EdU) staining, cells were pulsed with 10 µM EdU 24 hours prior to fixation with 4% paraformaldehyde. EdU incorporation was detected according to the manufacturer’s protocol using Click-iT^TM^ EdU Alexa 488 kit (Thermo Fisher, C10337). Alkaline Phosphatase (ALP) staining was performed on live cells using VECTOR Red ALP substrate kit (Vector Laboratories Cat#SK-5100) with a 30-minute incubation, according to the manufacturer’s instructions. The primary antibodies used in this study were as follows: Chromogranin A (CHGA, Abcam Cat#ab15160), Mucin 2 (MUC2, Santa Cruz Cat#sb-515032), Zonula Occludins-1 (ZO-1, Proteintech Cat#66452-1-Ig), or E-cadherin (Proteintech Cat#20874-1-AP). Secondary antibodies were as follows: Alexa Fluor™ 594 Goat Anti-Rabbit antibody (Jackson ImmunoResearch, Cat#111-585-003), Alexa Fluor™ 488 Goat Anti-Mouse antibody (Jackson ImmunoResearch, Cat#111-545-003), or Alexa Fluor™ 647 Goat Anti-Mouse antibody (Invitrogen, A21236). For immunocytochemistry staining, cells were fixed in cold 100% methanol (ZO-1 and E-Cadherin) or in 4% paraformaldehyde (MUC-2 and CHGA) for 30 minutes. Fixed cells were permeabilized using 0.5% Triton X-100 (Promega, Cat#H5142). Primary antibodies were added at 1:250 dilution in 1X Animal-Free Blocking Solution (Cell Signaling Technology, 15019L) to the apical side of each Transwell^®^ for an overnight incubation at 4°C. Secondary antibodies and Hoechst 33342 Stain (Invitrogen, Cat#H3570) were diluted 1:1,000 in 1X Animal-Free Blocking Solution and incubated on the apical side of each Transwell^®^ for 1 hour.

### Imaging

EdU images were acquired with the 10x objective lens using the ImageXpress^®^ Nano Automated Imaging System with MetaXpress Software version 6.5.4.532 (Molecular Devices). The post-laser offset and exposure time were adjusted to acquire a focus point and intensity that was comparable across the plate. Images of cell lineage and tight junction staining were acquired on an Olympus IX2-UCB microscope (Olympus, Shinjuku City, Tokyo, Japan).

### Histology

Hemotoxylin and Eosin (H&E) and Alcian Blue/Periodic Acid Schiff (AB/PAS) staining were performed on sections from 12-well RepliGut^®^ Planar cultures fixed in 4% paraformaldehyde on day 3 in RMM, according to the methods above. Fixed tissues were processed on a Leica ASP 6025 tissue processor, embedded in paraffin (Leica Paraplast), and sectioned at 5 µm thickness. Tissue sections were baked at 60°C for 60 minutes, deparaffinized in xylene, hydrated with graded ethanols, and stained with H&E or AB/PAS using a Leica Autostainer XL. For H&E, slides were stained with Hematoxylin (Richard-Allen Scientific, 7211) for 2 minutes and Eosin -Y (Richard-Allen Scientific, 7111) for 1 minute. Clarifier 2 (7402) and Bluing (7111) solutions from Richard-Allen Scientific were used to differentiate the reaction. For AB/PAS, the slides were stained with Alcian Blue (Anatech, LTD, 867) for 10 minutes, immersed in Periodic Acid (ThermoFisher Scientific, A223-100) for 5 minutes, rinsed in water, then transferred to Schiff reagent (Fisher Scientific, SS32-500) for 30 minutes followed by a Sulfurous rinse for 1 minute, and washed in running tap water for 10 minutes. Histology images were captured using the 40x objective on a Leica DMi8 microscope with an Amscope 18MP Aptina Color CMOS camera and AmScope software (Version 4.11.18421).

### Gene expression

For gene expression profiling, proliferative cells were collected at 3 days post-plating in RGM media and differentiated cells were collected 3 days after switching cultures to RMM media. At the time of collection, cells were rinsed once with 1X phosphate-buffered saline (PBS) and then collected in 500 µL per Transwell^®^ of RNA Lysis Buffer from the Ambion RNAqueous kit (Invitrogen AM1912). The Ambion RNAqueous kit was used to isolate RNA based on manufacturer’s protocol. RNA concentration was quantified via Qubit. Reverse transcription was performed using the iScript™ cDNA Synthesis Kit (BioRad, 1708891). Gene expression analysis using the Biomark HD qPCR System and Dynamic Array IFCs for Gene Expression (Fluidigm) was performed at the Advanced Analytics Core Facility at the University of North Carolina at Chapel Hill School of Medicine. Taqman^®^ probe Assay IDs for all genes analyzed are found in Table S1. Relative gene expression (2^−ΔΔ*Ct*^) for each gene was calculated by comparing the sample to the average proliferative cells value from that donor.

In the cell passage number analysis, Ct values were converted to ΔCt using 18S as the housekeeping gene. All subsequent analysis included the 91 remaining genes (i.e., all genes except 18S). Principal component analysis, hierarchical cluster analysis, and inter-replicate correlation analysis were performed to assess sample quality and agreement between replicates in each passage. Corresponding quality control (QC) plots were generated and used to determine outlier samples and assess similarity between each passage group. Heatmaps of ΔCt values for the set of 91 genes (including housekeeping genes GAPDH and ACTB) were generated to confirm the findings from the QC analysis. Outliers were removed, and all analysis and plotting were re-performed to assess the final set of samples. Differential expression analysis was performed using a generalized linear model (GLM) with the passage number as a factor. This analysis used linear regression to compare all the passages. Foldchange, p-value, and adjusted p-value (Benjamini and Hochberg, 1995) were calculated. Differentially expressed genes (DEGs) were identified for the comparison of each higher passage (5, 10, or 15) with passage 2. To be considered a DEG, a gene had to meet the following criteria: Absolute foldchange ≥ 1.5 and Adjusted p-value ≤ 0.05.

### LDH Cytotoxicity Assay

Cytokine-induced cytotoxicity was measured using the CyQUANT™ LDH Cytotoxicity Assay (Invitrogen, C20300). Supernatant was collected from both apical and basal compartments at 48 hours post treatment and pooled in a representative ratio of total media (25 µL apical and 50 µL basal). LDH activity was determined on the combined media following the manufacturer’s protocol. Absorbance was measured at 490 nm and 680 nm using a BioTek Synergy H1 plate reader. The absorbance of the media blank was subtracted from each sample, followed by subtracting the absorbance at 680 nm (background) from the absorbance at 490 nm. Lysed cells served as a maximum LDH release control. “% Cytotoxicity” was calculated using the following formula:

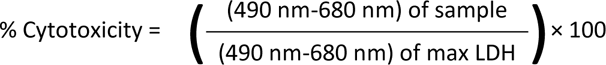

Dose response curves were generated using three parameter nonlinear regression. R-squared values comparing TEER to cytotoxicity were determined using nonlinear best fit line.

### Enzyme-Linked Immunosorbent Assays (ELISAs)

Interleukin 8 (IL-8) and C-X-C motif chemokine 11 (CXCL11) ELISAs were performed using basal supernatants collected 48 hours after cytokine treatments using IL-8 Human ELISA Kit (ThermoFisher, Cat# KHC0081) and Human CXCL11/I-TAC Quantikine ELISA Kit (R&D Systems, Cat# DCX110), respectively, following the manufacturer’s protocol. Absorbance at 450 nm was measured using a BioTek Synergy H1 plate reader. Background absorbance was subtracted from all data points, including standards, samples, and controls, prior to plotting. A standard curve was generated using a sigmoidal four parameter algorithm on BioTek software from which concentrations of detected proteins were calculated.

### Statistical Analyses

All statistical tests were performed in GraphPad Prism 9 (GraphPad, CA, USA). Statistical significance was set at a *P* value of <0.05 for all analyses. Kruskal-Wallis test was used to identify significant differences in TEER metrics between cell passage numbers. One-way ANOVA with Tukey’s multiple comparison test was used to test for significance between cytokine treatments with and without adalimumab and tofacitinib.

## 3 Results

### Sequential proliferation and differentiation over the RepliGut^®^ Planar culture lifespan

Transverse colon intestinal epithelial cells were derived from transplant grade donors and do not have a disease status. Demographics of the donors used in this study are shown in Table 1. A schematic of the RepliGut^®^ Planar model with the time-course of development and representative TEER is shown in Figure 1. Crypt-resident intestinal proliferating cells were first plated at sub-confluence onto hydrogel-coated transwell membrane inserts. Cells are grown to confluence using RepliGut^®^ Growth Medium (RGM) formulated to promote cell proliferation. Once cells reach confluence (4-6 days), RGM is removed and replaced with RMM to promote cellular polarization and differentiation into post-mitotic lineages (Figure 1A) (Gracz and Magness, 2014). Confluence is initially observed via brightfield microscopy and then confirmed by an increase in TEER above 250 Ω·cm^2^, at which point cell media is changed to RMM (Figure 1B). TEER continues to increase for about 2 days followed by a 3-5 day plateau phase (Figure 1B). The TEER plateau begins on the day when the average TEER of the 96-well plate meets or exceeds 75% of peak TEER and continues until TEER drops below 60% of peak and/or the coefficient of variation (%CV) between wells exceeds 25%. Similar to *in vivo* physiology, intestinal epithelium in cell culture has a finite lifespan once fully differentiated, lasting 3-10 days after differentiation begins (Snippert et al., 2010).

**Figure 1.**
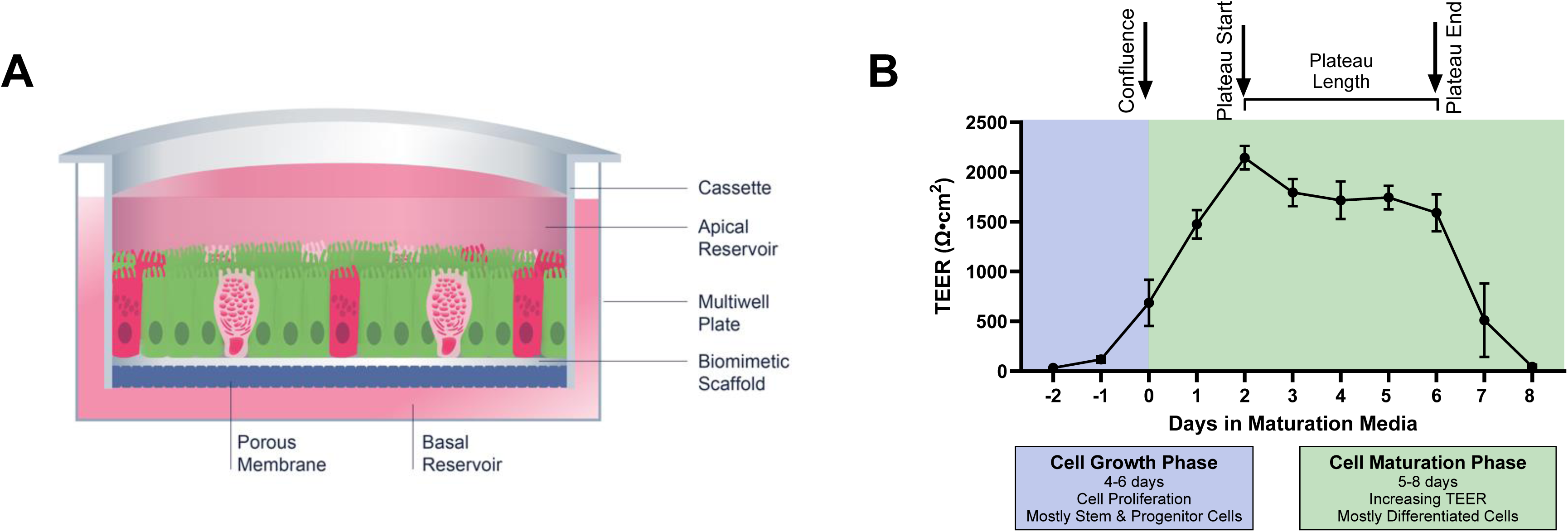
Transverse colon RepliGut^®^ Planar Model. (A) Cross section schematic of the RepliGut^®^ Planar Transverse Colon model comprised of multiple epithelial cell lineages. Cells derived from transverse colon stem and progenitor cell populations are seeded on a Transwell^®^ cassette coated with a biomimetic scaffold and provided with differentiation cues post-confluence. (B) Representative TEER curve of RepliGut^®^ Planar Transverse Colon over culture time course. The relative timeframe of each key measurement, time to confluence, time to plateau, and plateau length, that are used to characterize barrier formation and integrity of the cell monolayer are denoted above the graph. Data are represented as mean ± SD.

**Table 1.** Donor Characteristics.

Table and figure legends
| Table 1. Donor Characteristics |  |  |  |  |  |  |
| --- | --- | --- | --- | --- | --- | --- |
| Donor | Age (years) | Sex | Race | Height (in) | Weight (lbs) | BMI |
| 1 | 23 | Male | Caucasian | 75 | 182.1 | 22.8 |
| 4 | 51 | Female | Caucasian | 59 | 158.4 | 32.0 |
| 5 | 50 | Male | African American | 70 | 162.0 | 22.7 |
| 6 | 51 | Male | African American | 60 | 219.0 | 31.3 |

A unique characteristic of the RepliGut^®^ Planar model is the ability to induce the transition from a proliferative cell state to a differentiated confluent epithelium sequentially over the course of 10-12 days in culture. To characterize the dynamic lifespan of the RepliGut^®^ Planar model, markers of cell proliferation and differentiation at multiple time points in culture were assessed via microscopy and gene expression. Cells were pulsed with EdU for 24 hours to assess the proliferative capacity on days 1 through 8 of culture (Figure 2A) with a change from RGM to RMM on day 4 corresponding to observed confluence. Representative micrographs show positive EdU staining on days 2-4 in culture. On days 5 and 8, EdU staining was not detected, demonstrating that cells lost their proliferative capacity after the addition of RMM on day 4. Qualitative assessment of differentiated cell markers revealed the presence of absorptive and secretory cell lineages, as seen through positive staining of ALP, MUC2, and CHGA (Figure 2B). E-cadherin and ZO-1 staining of differentiated cells also indicated the formation of tight junctions between days 4-6 in culture (Figure 2B).

**Figure 2.**
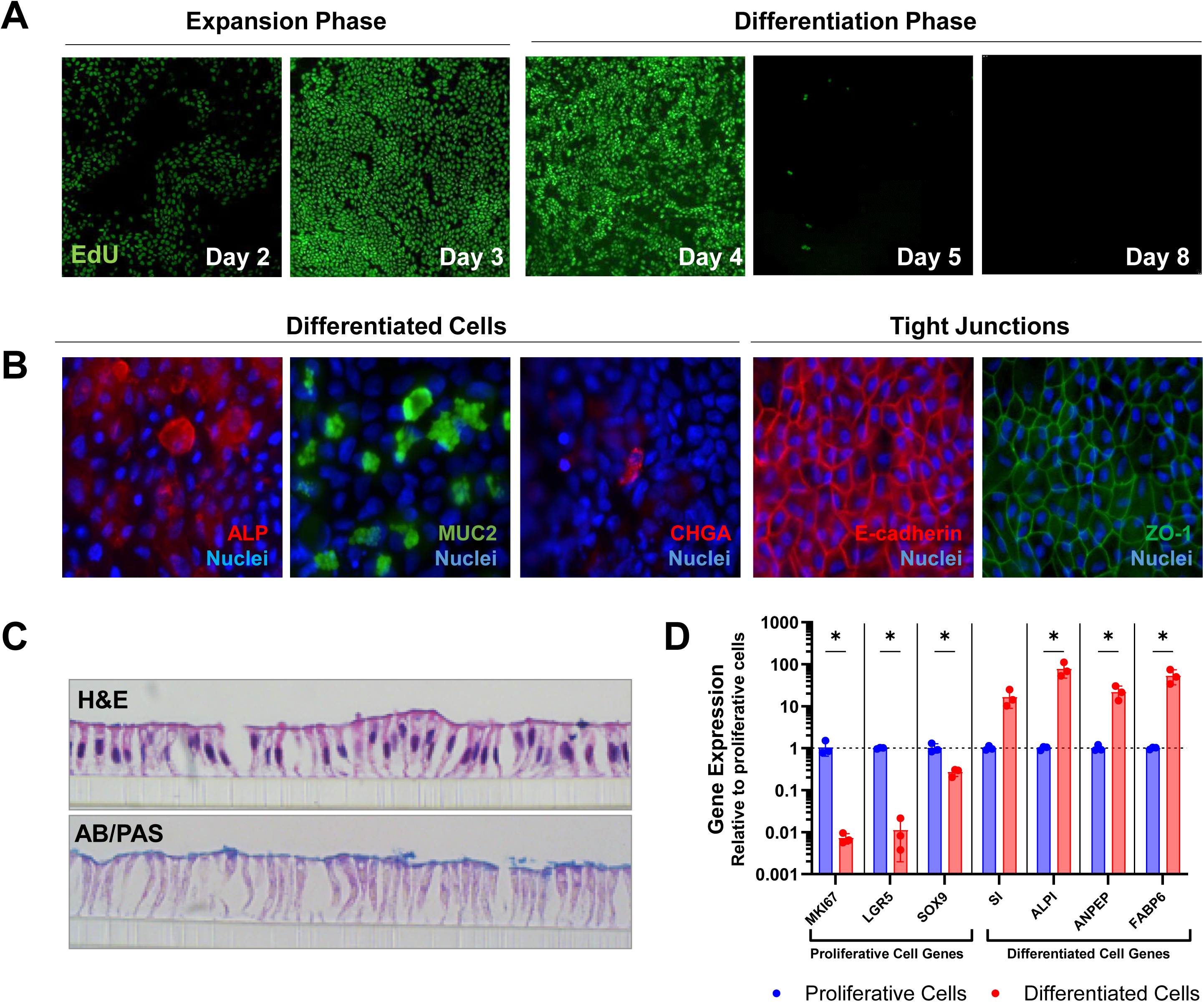
Phase specific model characteristics. (A) Representative images of EdU incorporation over time. (B) Representative images of fully differentiated cells stained for cell proteins ALP (red), MUC2 (green), and CHGA (red), DAPI (blue) and tight junction proteins ZO-1 (green) and E-cadherin (red). (C) Representative histology image of monolayer cross sections in the differentiated phase stained for H&E (top) and for AB/PAS (bottom). (D) Gene expression of proliferative and differentiated cell genes in cells in proliferative and differentiation phases. 2^ΔΔCt^ values were calculated, normalized to a 18S housekeeping gene and reported relative to proliferative phase cells. *p< 0.05, t-test with Welch’s correction. Data are represented as mean ± SD.

To confirm that cells had polarized after the addition of RMM, we evaluated hematoxylin and eosin (H&E) staining of histological sections of cells derived from one human donor (Donor 5) on day 3 in RMM. H&E staining revealed a polarized columnar epithelium (Figure 2C). We further assessed the presence of mucopolysaccharides via AB/PAS staining. Positive staining was localized to the apical surface, indicating the presence of mucus-producing cells with correct directional secretion (Figure 2C), further confirming the differentiated and polarized cell physiology.

To support these morphological observations, we assessed expression of genes associated with cellular proliferation and differentiation from Donor 5 on Day 3 post-plating (proliferative phase), and Day 3 in RMM (differentiated phase) in culture corresponding to the days of highest and lowest EdU incorporation. Relative to proliferative cells, differentiated cells had lower expression of genes associated with proliferation (MKI67, LGR5 and SOX9), and higher expression of genes associated with differentiation (SI, ALPI, ANPEP and FAPB6) (Figure 2D). All differences in gene expression between proliferative cells were significant except for SI (t-test with Welch’s correction, p<0.05). Taken together, these data indicate that the RepliGut^®^ Planar model is comprised of cells with transient proliferative capacity that can differentiate into multiple cell lineages, mimicking the cellular morphology of human intestinal epithelial cells (Gracz and Magness, 2014; Bhatt et al., 2018; Rees et al., 2020).

### Cell passage number is a source of variability in the RepliGut^®^ Planar model

Cell passage number has been linked to variability in Caco-2 cell culture, causing effects on gene expression, protein production and overall cell function (Sambuy *et al*., 2005). Somatic stem cells such as those in the intestinal crypt may have a finite ability to self-renew and differentiate appropriately which may be accelerated in a cell culture environment (Snippert *et al*., 2010; Liu and Rando, 2011). We therefore evaluated whether cell passage number could be a source of variability in the RepliGut^®^ Planar model. Gene expression analysis was performed using qPCR and included 91 genes corresponding to known identity and functional markers of intestinal epithelium, including enterocyte, secretory, metabolism, and inflammatory signaling related genes (Table S1). Genes were analyzed relative to passage 2, which is the earliest cell passage possible after the initial freeze. Principal component analysis, hierarchical cluster analysis, and inter-replicate correlation analysis of ΔCt values revealed that passage 15 samples separated from the rest of the passages (Figure 3A, 3B). Differential expression analysis performed using a generalized linear model identified 29 DEGs between p15 vs p2, 2 DEGs between p10 vs p2, and no DEGs between p5 vs p2 (Fig 3C). This analysis suggests that p10 is the highest passage that resembles the native cells (p2) of the passages we tested.

**Figure 3.**
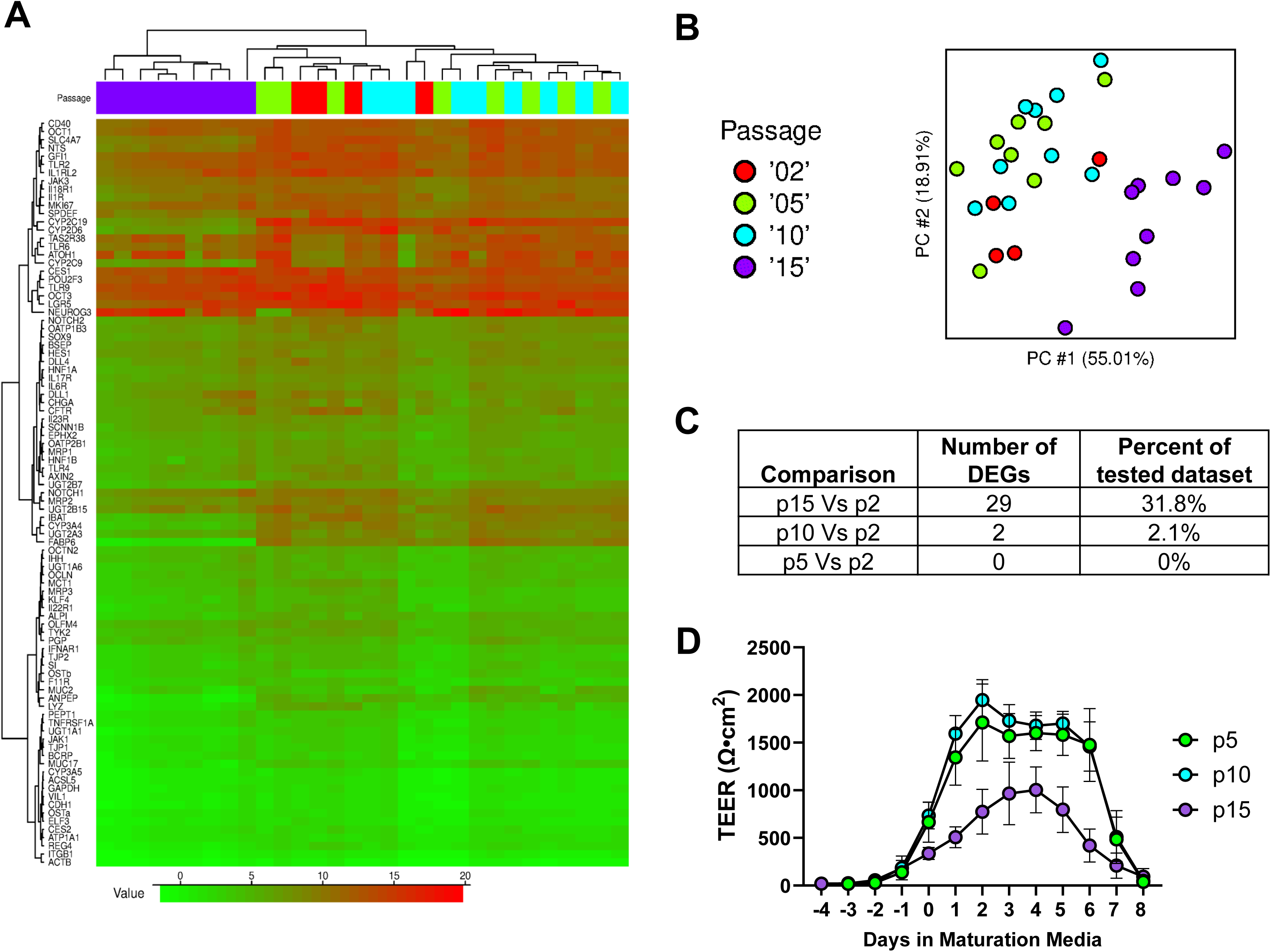
Cell passage number is a source of variability. (A) Hierarchical cluster analysis heatmap of the 91 tested genes (by ΔCt value) of passage 2, 5, 10 and 15 cells. (B) Principal component analysis of the different cell passage numbers. (C) Number of differentially expressed genes of each cell passage number relative to passage 2 cells. (D) TEER of the different cell passage numbers. Data are represented as mean ± SD.

To determine whether gene expression patterns among different cell passage numbers related to functional and morphological differences during differentiation, we measured barrier integrity daily using TEER until day 8 in RMM from p5, p10, and p15-derived models during maturation phase. Having shown that p5 has relatively few gene expression differences compared to p2, p2 cells were not included in this analysis. TEER curves from four independent runs were generated and averaged for each cell passage to observe time-driven variability in barrier formation and integrity over the duration of the RepliGut^®^ Planar culture (Figure 3D, Figure S1). No significant differences were observed in time to confluence (p>0.05, Kruskal-Wallis test). P15 cells reached significantly lower peak TEER than p10 cells (p=0.04, Kruskal-Wallis test; Figure 3C-F). By day 2 in RMM, p5 cells reached a peak TEER of 1826 ±366 Ω·cm^2^ and p10 cells reached a peak TEER of 1949 ±169 Ω·cm^2^, whereas p15 cells only reached a peak TEER of 1032 ±324 Ω·cm^2^ by day 4 in RMM (Figure S1). Relative to p15 cells, p10 and p5 cells had a longer TEER plateau length with a significant difference between p10 and p15 (p=0.0381, Kruskal-Wallis test). The median plateau lengths were 4.5, 5, and 3 days for p5, p10 and p15 cells, respectively (Figure S1). Interestingly, these plateau lengths are representative of the expected 3-5-day life span of intestinal epithelial cells (Reynolds *et al*., 2014; Rees *et al*., 2020). This analysis demonstrated that while passage number does alter the magnitude of peak TEER and overall shape of the TEER profile, it does not meaningfully affect the ability of cells to form or maintain a confluent monolayer with tight junctions.

Together, these data suggest a fundamental change to the stem-cell identity that translates into functional differences of the epithelial monolayer and temporal behavior of the model. To avoid the possibility of stem cell identity drift affecting model performance, cells beyond passage 10 were not used in subsequent analyses.

### Barrier formation and maintenance metrics are consistent across donors

Having determined that p10 cells resulted in cultures with similar barrier formation kinetics and gene expression patterns to p2 cultures, we moved forward to characterize donor-donor, lot-lot, and run-run variability in p10 cultures from 18 independent cell production lots across 4 donors (n = 5, 3, 9, and 1 cell lots for Donors 1, 4, 5, and 6, respectively). Donors 2 and 3 have yet to be characterized and were excluded from analysis. Post-thaw viability of ≥ 75% [87 ± 6%] was observed for all cell lots generated (Figure 4A), with no significant differences between Donors (Kruskal-Wallis test). Time to achieve culture confluence (Figure 4B) ranged from 2 to 9 days, with medians of 4, 4, 4, and 5 days for Donors 1, 4, 5, and 6, respectively. No significant differences were observed between donors in time to confluence (Kruskal-Wallis test).

**Figure 4.**
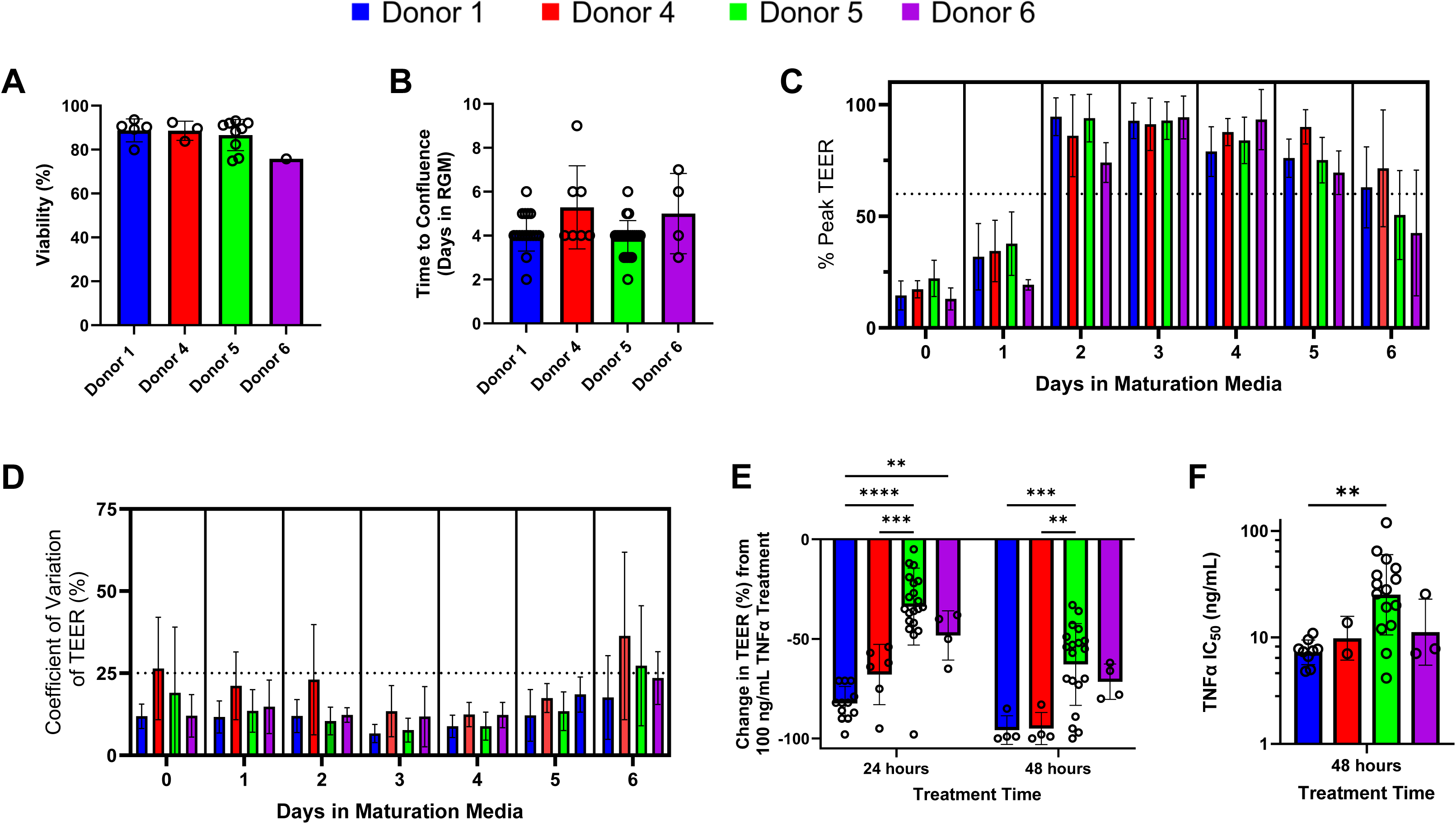
Consistency of TEER curve metrics between donors. (A) Bar graph showing the viability post thaw for the cell lots within each donor (each open circle representing independently generated lots of p9 stem cells). (B-D) 51 independent model runs consisting of lots/runs for each donor as follows: Donor 1: 5/17, Donor 4: 3/7, Donor 5: 9/23, Donor 6: 1/4 evaluated for: the time to confluence measured in days (B), % of peak TEER for each day in RMM with dotted line at 60% (C); and % CV for TEER values at days in RMM with dotted line drawn at 25% CV (D). Each bar represents CV from 24 wells for each of the 51 total runs. (E) Time-dependent and donor differences in the effect of 100 ng/mL TNFα on TEER, % change normalized to vehicle control. (F) Time-dependent and donor differences in TEER sensitivity to TNFα. Bar graph comparing the average IC-50 for TEER values determined from -12 doses of TNFα starting day 2 in RMM and continuing for 24 or 48 hours as indicated for each donor.

Essential to the utility of the culture model is the predictable formation and maintenance of barrier integrity to provide a reasonable assay window. To characterize barrier kinetics and variability, TEER was measured across multiple days and multiple experiments using the 18 lots across four Donors described above, representing a total of 51 experiments. To allow comparison across lots and experiments, maximum TEER within each experiment was determined and all other TEER values were normalized to the within-run peak TEER. Out of the 51 experiments included in this analysis, only four runs did not reach 75% of peak TEER by Day 2 in RMM. Using a within run acceptable variability cutoff of CV<25%, only two experiments (Figure 4D) exceeded the acceptable level on Day 2 in RMM (Donor 4, one of which carried on to Day 3), and only one experiment exceeded the acceptable level on Day 5 in RMM (Donor 1). Together, this data supports predictable culture behavior with lowest experimental variability between Days 3 and 5 in maturation media, providing a clear 72-hour assay window during TEER plateau that enables meaningful assessment of barrier function using TEER. Figure 4D also includes within-plate variability data for TEER at Day 0 in RMM. Many cultures at this timepoint still demonstrated some degree of variability in TEER. We suspect this to correspond to some instances of spontaneous differentiation; as cultures mature, within-plate variability in TEER diminishes.

The cytokine signaling molecule TNFα plays a significant role in the etiology of inflammatory bowel diseases. In addition to cytotoxicity, these proinflammatory cytokines promote epithelial cell-chemokine release to recruit and activate immune cells involved with tissue damage and repair (Dwinell *et al*., 2001; Kucharzik *et al*., 2005; Treede *et al*., 2009; Sonnier *et al*., 2010; Friedrich, Pohin and Powrie, 2019). To investigate whether donors have a variable response to TNFα, we analyzed TEER reduction at 24 and 48 hours post-TNFα treatment in multiple cell lots from all four donors. At 24 hours post-treatment, Donors 1 and 4 had greater reductions in TEER compared to Donors 5 and 6, which increased to comparable levels by 48 hours (Figure 4E). At 24 hours post-treatment, significant differences were detected between Donors 1 and 5, Donors 1 and 6, and Donors 4 and 5 (Two-way ANOVA with Tukey’s multiple comparisons test). At 48 hours post-treatment, significant differences were detected between Donors 1 and 5, and Donors 4 and 5 (Two-way ANOVA with Tukey’s multiple comparisons test). Data shown is from 43 of 51 experiments represented in Figure 4 C & D; not all experiments included TNFα dosing. Additionally, 9 data points (six from Donor 1 and two from Donor 4) were removed from the 48 hour data sets due to a smaller percent change in TEER observed as compared to the same wells at 24 hours, a phenomenon believed to result from excess cell debris following cell death. One data point from Donor 5 was removed from all TNFα analyses due to a lack of TEER response at any dose tested, believed to result from a technical error. To further characterize donor-to-donor variability, the 48 hour IC_50_s for percent change in TEER were compiled (Figure 4F). The IC_50_s and 95% confidence intervals for Donors 1, 4, 5, and 6 were 7 [6, 8], 10 [8, 12], 25 [24, 27] and 11 [9, 13] ng/mL, respectively. The 95% confidence intervals on the IC_50_ reflect low variability; only for a single donor, 5, did the range of all observed IC_50_s exceed one log. A significant difference in IC_50_ was only observed between Donors 1 and 5 (Kruskal-Wallis, p=0.0062). Together, these data suggest an inherent variability in biological sensitivity to proinflammatory insult that increases over time.

### Human donors exhibit varying responses to proinflammatory cytokines

Having identified culture conditions where baseline lot to lot and run to run variability are minimized across donors, we tested functional responses to the pro-inflammatory cytokines TNFα and IFNγ that are commonly involved in manifestation of IBD (Andreou, Legaki and Gazouli, 2020; Gareb *et al*., 2020). Increasing concentrations of TNFα or IFNγ were applied to RepliGut^®^ Planar cultures derived from four donors on day 2 of differentiation and TEER was monitored over 48 hours, at which point cells were analyzed for cytotoxicity via LDH release and for IL-8 or CXCL11 chemokine release. All donors elicited a dose-dependent response characterized by TEER reduction and LDH increase to both TNFα and IFNγ treatments with concomitant associated increase in IL-8 or CXCL11 release, respectively (Figure 5A-C, Figure S2).

**Figure 5.**
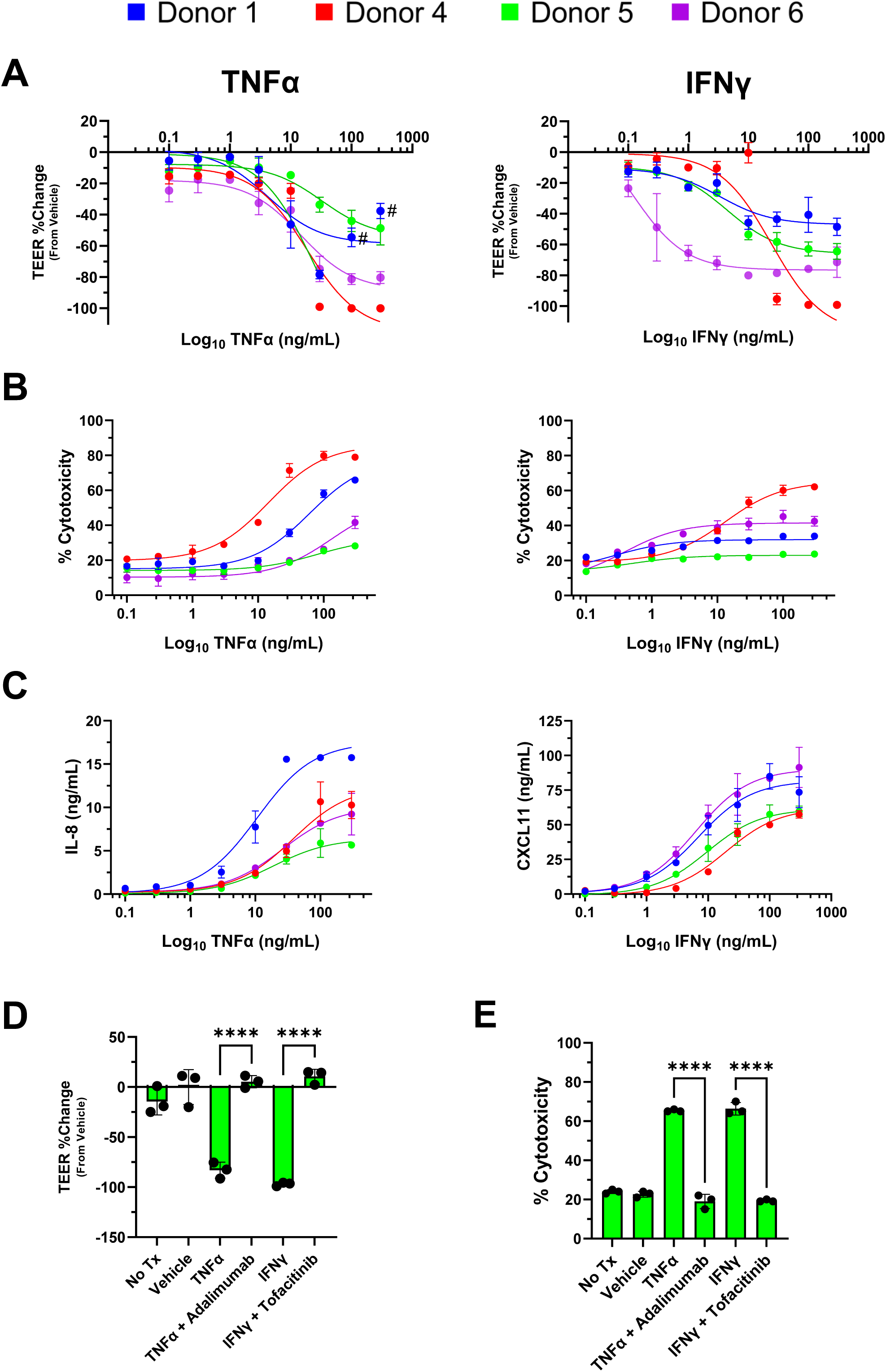
Donor-to-donor differences in TNFα and IFNγ response. (A) Percent change in TEER relative to vehicle after 48 hour treatment with TNFα (left) or IFNγ (right) of four donors. (B) Percent cytotoxicity relative to a maximum LDH activity treatment. (C) IL-8 (left) or CXCL11 (right) release in response to increasing doses of TNFα (left) or IFNγ (right). (D-E) Percent change in TEER and percent cytotoxicity of cells treated with 300 ng/mL TNFα or 300 ng/mL IFNγ in the presence or absence of 3 µg/mL of adalimumab or 100 μM tofacitinib for 48 hours. ****p<0.0001, One-way ANOVA with Tukey’s multiple comparison test. Data are represented as mean ± SD. n = 3.

IC_50_ values for TEER reduction in response to TNFα were comparable between donors (14.68-32.8 ng/mL), demonstrating low donor-to-donor variability in sensitivity to TNFα, though the magnitude in TEER reduction was greater for Donor 1 (Figure 5A, left panel). After 48 hours of treatment with the two highest doses of TNFα, increased TEER relative to the maximum response in Donor 1 was observed, consistent with excessive cells debris that clogs the pores of the membrane. Due to increased interference from cell debris, we excluded two dose points from TEER IC_50_ calculations (shown as annotated points in Figure 5A, left panel). In contrast, sensitivity to IFNγ was variable across Donors, with Donor 6 observed to have the highest sensitivity to IFNγ (IC_50_ 0.12 ng/mL), compared to the other three Donors. The magnitude of response to IFNγ varied by 2.5-fold across the four Donors with Donor 1 showing the smallest reduction in TEER (-48.52±5.64%) and Donor 4 exhibiting the greatest reduction in TEER (-99.18±0.122%) (Figure 5A, right panel).

All four donors displayed a dose dependent increase in cytotoxicity in response to TNFα (Figure 5B, left panel). Similar to TEER kinetics, the magnitude of cytotoxicity in response to each cytokine was donor-dependent and correlated with barrier disruption across all four donors (R^2^ > 0.7) (Figure S3). Cytotoxicity responses to IFNγ were not as robust as TNFα responses (Figure 5B, right panel). To measure effects of TNFα and IFNγ on chemokine signaling pathways associated with NF-κB or JAK/STAT signaling, we measured release of IL-8 and CXCL11, respectively. In line with maximum response to TNFα mediated barrier disruption, Donor 1 elicited the highest levels of IL-8 release compared to Donors 4, 5 and 6 (Figure 5C). The concentration of CXCL11, a JAK/STAT regulated cytokine, increased similarly in cells treated with IFNγ in all four donors (Figure 5C). Together, these data reveal donor dependent sensitivities and maximum responses to proinflammatory cytokines.

To demonstrate that clinically validated TNFα- and IFNγ antagonists can protect against cytokine-induced barrier disruption and cytotoxicity, we co-treated cells from a single donor, Donor 5, with canonical marketed inhibitors of each cytokine, adalimumab or tofacitinib (Al-Bawardy, Shivashankar and Proctor, 2021; Antunes *et al*., 2021; Cai, Wang and Li, 2021), respectively. For this study, doses of TNFα and IFNγ were used that corresponded to the maximum response in barrier disruption and cytotoxicity in RepliGut^®^ Planar. Both FDA-approved clinical inhibitors effectively preserved barrier integrity (Figure 5D) and prevented cellular cytotoxicity (Figure 5E) induced by TNFα or IFNγ. These data demonstrate the utility of the RepliGut^®^ Planar as a tool to explore drug pharmacology associated with TNFα and IFNγ pathways in IBD.

## 4 Discussion

While spherical organoid models have provided an important opportunity to understand development of the intestinal epithelium, they have limited ability to model key functions of the epithelial barrier, owing to their spherical shape and fully-interior lumen (Wang *et al*., 2017; Dutton *et al*., 2019; Franco, Da Silva and Cristofoletti, 2021). The RepliGut^®^ Planar system is a 2D monolayer of intestinal epithelial cells derived from primary human intestinal resident crypt stem cells. This allows for technical access to both the basal and luminal sides of a polarized epithelial cell layer and overcomes limitations of 3D model systems. RepliGut^®^ Planar is established by inducing differentiation of a stem cell-derived proliferative cell population. As stem cell derived models are in a constant state of functional flux throughout culture duration, identifying appropriate conditions from which to build meaningful assays requires understanding of the factors driving variability in a time-dependent manner.

Initial study of gene expression patterns over time allowed us to verify proliferation and differentiation status through time in response to growth factor addition and removal (Figure 2D). Combined with functional evaluation of barrier formation and maintenance, the culture timeline can be separated into three phases (time to confluence, time to TEER plateau, and plateau length) to characterize the impact of donor, cell lot, and passage number variation on overall performance of the model (Figure 1B). Such evaluations enabled development of a robust assay within the TEER plateau phase that exhibited broad dynamic range for detecting barrier disruption and release of inflammatory chemokines, modeling key mechanistic events in inflammatory bowel diseases.

The observed TEER plateau lengths for p5 and p10 cells coincide with the expected 3-5-day life span of fully-differentiated intestinal absorptive epithelial cells that is observed *in vivo* (Reynolds *et al*., 2014; Rees *et al*., 2020). Together, these data imply that a shift in stem cell identity occurs after serial passaging original source cells in culture beyond p10, which may warrant further exploration. Analysis of four independent runs with cells from p5, p10 or p15 demonstrated that passage number does not limit functional barrier formation, as assessed via TEER (Figure 3B), though peak TEER values were much lower for p15 cells than p5 or p10. Alternatively, the time to plateau had considerable variability between p15 and p5. p5 and p10 cells reached peak TEER by day 2 in RMM, and plateau lengths averaged 4.5±0.57 days and 4.75±0.50 days, respectively.

Surprisingly, donor-to-donor variation was not as significant as expected within time to confluence and time to TEER plateau, but was in line with prior publications that also observed low variability among 3D organoids derived from multiple human donors (Mohammadi *et al*., 2021). One potential reason for strong donor-to-donor correlation is that RepliGut^®^ Planar lacks an intestinal microbiome and immune cells, which are key variables between humans that can alter clinical outcomes (Khan *et al*., 2019). Generating an intestinal barrier without a microbiome opens the door to model host-microbe interactions in a controlled environment using specific microbial components.

Donor-to-donor differences were most evident in functional responses to proinflammatory cytokines. While we observed reduced barrier integrity as early as 24 hours, some donors required 48 hours before a reduction in TEER was observed (Figure 4E). These data emphasize the importance of understanding the inherent biological variability from donor-to-donor when designing experiments that require timelines specific for other downstream readouts, such as those associated with cytotoxicity or pathway signaling. Nevertheless, cells from all four donors tested in the RepliGut^®^ Planar model responded to proinflammatory cytokines with increases in cell cytotoxicity, disruption of barrier, and increased secretion of the NF-κB and JAK/STAT-regulated chemokines IL-8 and CXCL11 respectively, demonstrating the utility of RepliGut^®^ Planar as an *in vitro* model for inflammatory studies (Roda *et al*., 2010; Okamoto and Watanabe, 2016).

One of the main objectives of this work was to understand both the opportunities and limitations of assays using a dynamic stem cell-derived model system. We observed that variability in TEER measurements increased with culture time, suggesting that the best opportunity to obtain meaningful assay responses are within earlier time windows following change to RMM. In this regard, variation in TEER was lowest (CV < 25%) between donors and lots during days 2-5 of the differentiation phase, which encompasses the TEER plateau phase of the RepliGut^®^ Planar model time-course (Figure 4). Therefore, we identified this phase as the optimal window to detect meaningful and reproducible acute responses to proinflammatory cytokines. Treatment of the RepliGut^®^ Planar culture with either TNFα or IFNγ at day 2 in RMM for 48 hours produced a dose-dependent chemokine production and disruption of the epithelial barrier as measured with TEER and LDH activity. Further, we demonstrated successful blockade of cytokine-induced epithelial damage using the FDA-approved clinical therapies adalimumab (neutralizing TNFα antibody) and tofacitinib (STAT inhibitor). Together, these data highlight the ability to use RepliGut^®^ as a drug discovery tool for anti-inflammatory drugs targeting these pathways.

In summary, gene expression and culture metric analyses revealed that cell passage number is a source of variability in the stem-cell derived RepliGut^®^ Planar platform. Comparison of passage 10 cells from multiple human donors indicated that there is low donor-to-donor variability in barrier formation and integrity, but variation is evident in the magnitude of response to proinflammatory cytokines.

## Data availability statement

All data supporting results and conclusions are contained within the article or supplementary material.

## Conflict of interest

CMP, BZ, MKB, BEM, MC, CB, JMM, JAL, DCG, RL, WT, MKB and EMB are current or previous employees of Altis Biosystems, Inc. DP, MBM, DM, RS are employees of Sciome.

### Acknowledgements

The authors thank Leah Huntress, Jacob Coyne, Erin Dancy, Lauren Boone, Earnest Taylor, Reganne Lorichon, and Vassili Kouprianov for their technical contributions to this work. We thank Mia Evangelista in the Pathology Services Core (PSC) for expert technical assistance with Histopathology. The PSC is supported in part by an NCI Center Core Support Grant (P30CA016086). We thank Gabrielle Cannon at the CGIBD Advanced Analytics Core for her expertise with BioMark. The CGIBD Advanced Analytics Core is supported in part by National Institute of Diabetes and Digestive and Kidney Diseases Grant P30 DK034987. This research was also funded by National Center for Advancing Translational Sciences Grant#1 R43 TR004230 and National Institute of Diabetes and Digestive and Kidney Diseases Grant# 1 R43 DK130708

## Supplemental tables and figure legends

**Table S1. TaqMan probe accession IDs and Ct values used in gene expression analysis**

**Figure S1.**
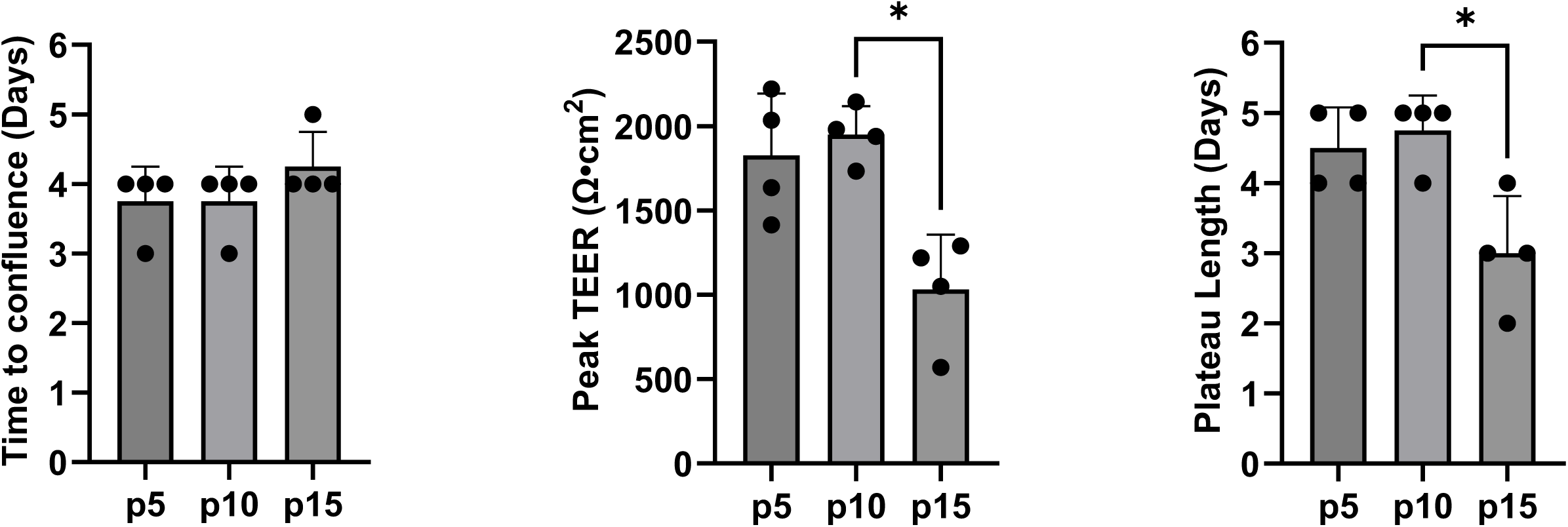
TEER metrics of different cell passage numbers. Bar graphs of four independent runs from each passage denoting days to confluence, peak TEER, and plateau length (*p<0.05, Kruskal-Wallis test). Data are represented as mean ± SD.

**Figure S2.**
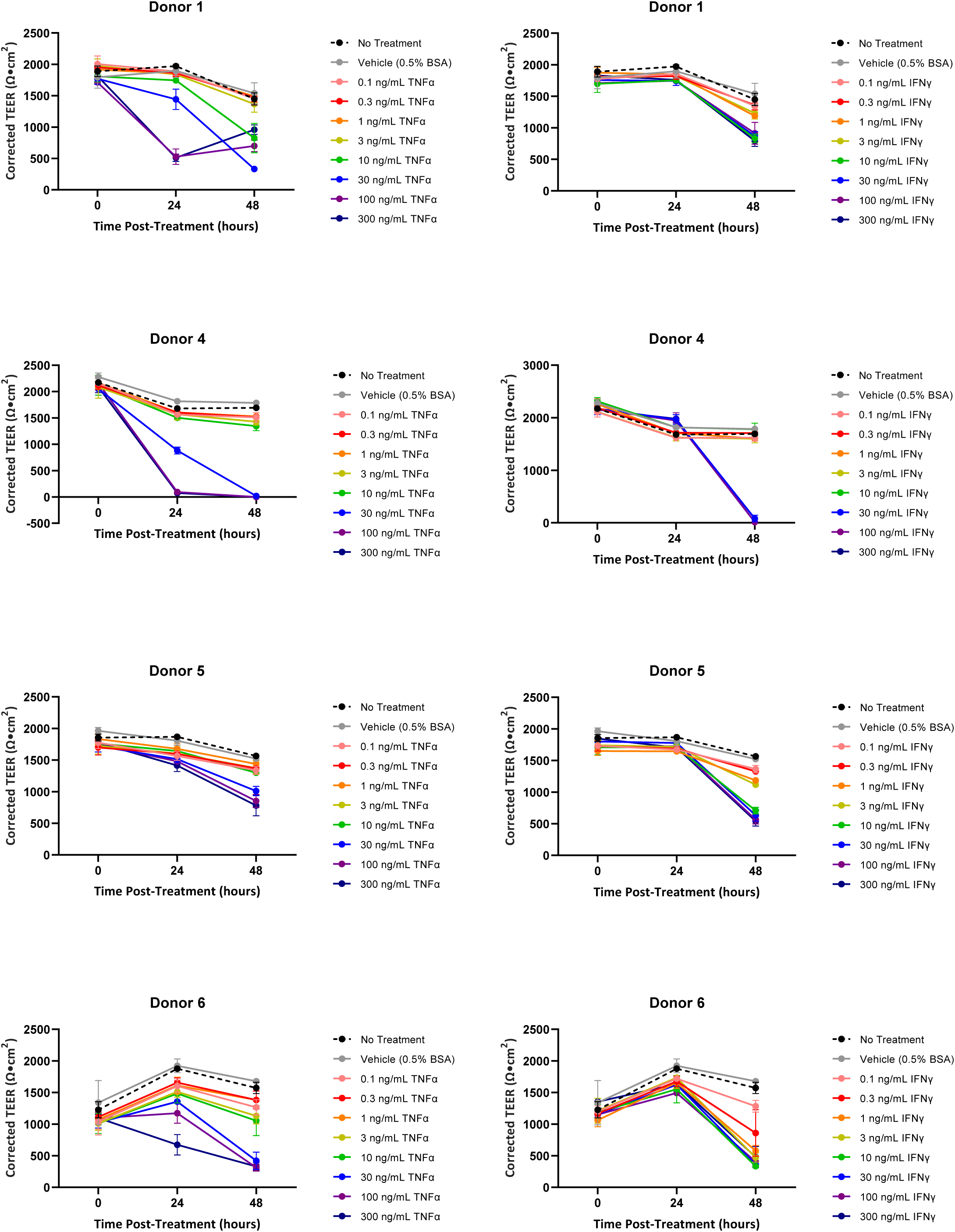
TEER of Donors 1, 4, 5 and 6 treated with TNFα or IFNγ from the RepliGut case study. Data are represented as mean ± SD.

**Figure S3.**
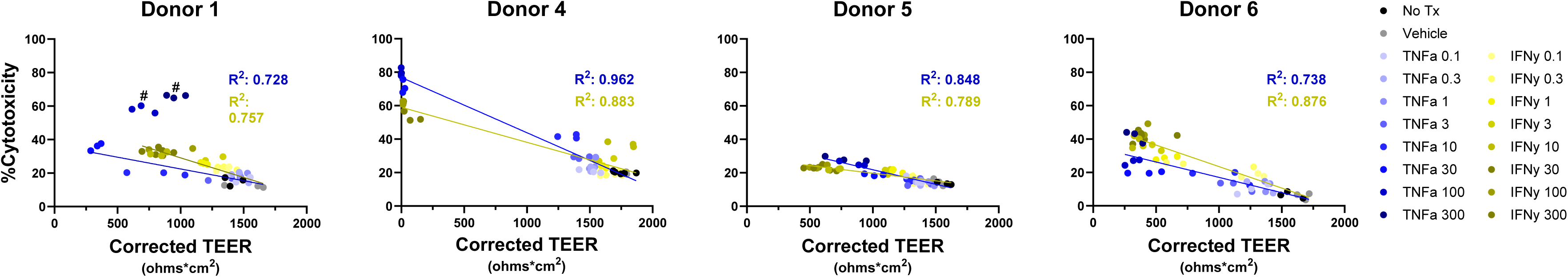
Donor-to-donor differences in LDH activity (% Cytotoxicity) in response to TNFα or IFNγ. Correlation plots between LDH activity increase and barrier decrease (TEER) for TNFα (blue) and IFNγ (yellow) for each donor as indicated. R^2^ values are noted in insets.

